# Stability of Commonly Used Hematological Parameters in Samples Stored at 33°C, 22°C and 4°C

**DOI:** 10.1101/148601

**Authors:** Ashish Jain, Sanchit Jain, Neha Singh, Priyanka Aswal, Shweta Pal, Sushant Kumar Meinia, Nilotpal Chowdhury

## Abstract

**Aim:** This study aimed to investigate the analytical bias and imprecision in haematological parameters induced by storage at 4°C, 22°C and 33 °C.

**Methods:** Three K_2_EDTA anticoagulated vials of blood were collected from each of twenty blood donors and stored at 4°C, 22°C and 33°C respectively. Readings from each vial were taken at 0, 4, 6, 12, 24, 48 and 72 hours after collection on the Sysmex XP-100 analyser. The mean and median shift of the parameters relative to the baseline and the coefficient of variation for each time-temperature combination were calculated. The shift was compared to the maximum acceptable bias.

**Results:** Haemoglobin, Red Blood Cell Count, White Blood Cell Count, Mean Corpuscular Haemoglobin were stable for at least twenty four hours at 33°C. Haematocrit, Mean Corpuscular Volume and Platelet Counts were stable for less than four hours at 33°C. All the above parameters were stable for longer at 22°C and 4°C. The three-part differential count showed instability within four hours at 33 °C.

**Conclusions:** Strict pre-analytical control is needed at 33°C or above due to the marked instability of most parameters. However, Haemoglobin, Red Blood Cell Count, White Blood Cell Count and Mean Corpuscular Haemoglobin remain relatively stable even at 33°C.

**Key Message:** Haematology samples exposed to temperatures of 33°C or above show rapid change in MCV, HCT,MCHC, RDW, Platelet Counts and three-part differential counts. Settings where prolonged exposure to these temperatures cannot be avoided should rely on the more stable parameters of Haemoglobin, RBC Counts, MCH and WBC Counts.

## Introduction

Knowledge of the effects of temperature on blood sample stability is important in a haematology laboratory and blood bank setting. Samples from peripheral collection centres and blood donation camps may have considerable delay in transport to the central laboratory. Errors in transport of vials cause further delays in sample processing. In such cases, the samples may be exposed to high temperatures, especially during summer and in tropical conditions. Such exposure of blood samples to high temperatures after collection is especially common in resource poor areas, but may also happen due to failure of temperature controlled transport and storage containers. Ideal conditions for storage are relatively uncommon in under resourced countries, many of which also have a warm to hot climate for a large portion of the year. A period of storage at high temperature may lead to variation in the different haematology parameters and adversely affect the accuracy of the final report. Therefore, it is important to know which parameters show clinically significant shift on storage under high temperatures, and which parameters remain relatively stable under the same conditions. This will aid in the proper interpretation of the haematology sample in case it becomes necessary to report on a sample known to have been exposed to high temperatures.

The analytical stability for the different haematological parameters has been studied by various researchers^1–6^. Most of these studies have examined the average bias on storage under standard conditions (around 4°C and room temperature). Very few studies have examined the stability of haematological samples at temperatures above 30°C, which are common in the tropical countries. Most previous studies also have not documented changes in analytical precision on storage.

The present study was designed to partly address the above limitations. In this study, the analytical stability of different haematological parameters over a period of seventy two hours in samples stored at 33°C, 22°C and 4°Cwere estimated. Thus, we attempted to examine the stability of the haematological parameters in blood samples on storage at temperatures likely to be encountered in many tropical countries as well as under ideal storage conditions.

## Subjects and Methods

The study was carried out in the Department of Transfusion Medicine and Blood Bank after obtaining ethical clearance from the institutional ethics committee. The study complied with the World Medical Association Declaration of Helsinki regarding ethical conduct of research involving human subjects.

In this study, samples collected from the diversion pouches of twenty apparently healthy (with no known comorbidities) blood donors were used. The blood donors were selected according to criteria set by regulatory agencies^7^. Additionally, donors with ingestion of antiplatelet drug were not included in the study.

Three samples, each of three ml, were collected from each donor in K_2_EDTA vials. Three baseline readings were taken from each of the three vials within 30 minutes of collection (thus having 9 baseline readings in total) on the Sysmex XP-100 three part analyser (Sysmex Corporation, Kobe, Japan). The readings of the following parameters were studied: White Blood Cell Count(WBC), Red Blood Cell Count(RBC), Haemoglobin(HGB), Haematocrit (HCT), Mean Corpuscular Volume (MCV), Mean Corpuscular Haemoglobin (MCH), Mean Corpuscular Haemoglobin Concentration (MCHC), Red Cell Distribution Width (RDW), Platelet Count (PLT), Mean Platelet Volume (MPV), and the Differential Count parameters, viz. the Lymphocyte Count(LC), Mixed Cell Count(MCC) and Neutrophil Count (NCC).The mean of the nine baseline readings of the above parameters were calculated and accepted as the baseline value for each parameter. The baseline within-subject coefficient of variation (CV) was also calculated. After the baseline readings, the samples were stored. Of the three samples collected from each donor, one was stored under refrigeration (4° C); the second was stored at 22°C and the third sample at 33°C.

These stored samples were analysed after four hours, six hours, twelve hours, twenty four hours, forty eight hours and seventy two hours of storage. At each of the above mentioned time points, three readings of all the parameters from each sample were taken, after which they were re-stored at their respective storage temperature. The mean of the three readings for each parameter of interest was calculated at each time point for each temperature, and accepted as the representative value of that parameter for that particular storage time at that particular temperature. The within subject coefficient of variation for each parameter of interest for each time point at each temperature was also calculated. The percentage difference of the representative value for each parameter (at each time point and temperature) from the baseline value was calculated. The flow chart of the methods followed up to statistical analysis is given in Figure 1. The flow chart of the statistical decision-making process is given in Figure 2.

**Figure 1:**
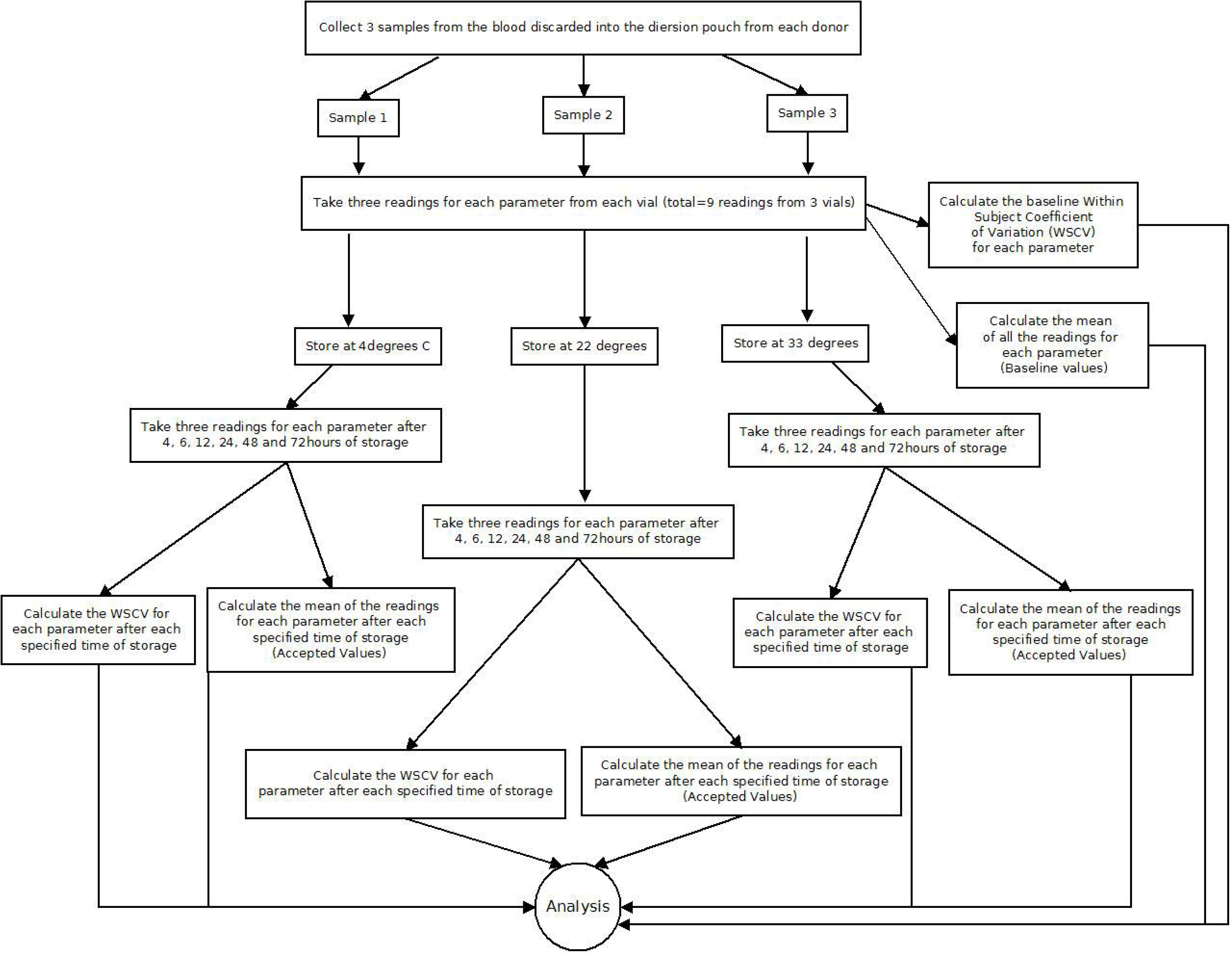
Flow chart showing the methods followed in the study up to the statistical decision making.

**Figure 2:**
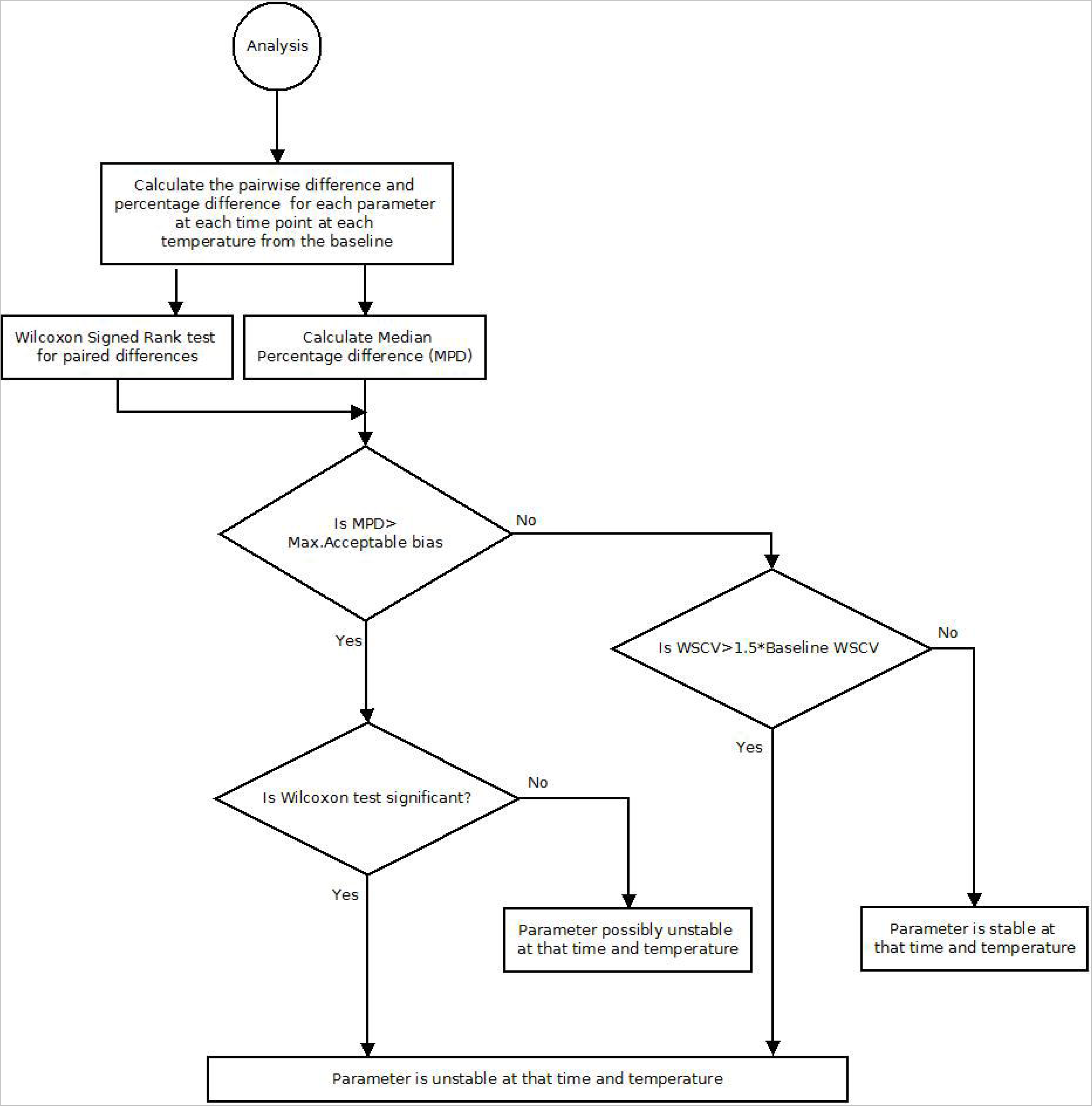
Flow chart of the decision making in the present study.

All the measurements were done under the Whole Blood Mode on Sysmex XP-100 three part analyser with manual feeding of sample. The instructions in the analyser instruction manual were strictly followed. The collection and processing of the samples were done by the same person. Quality control of the analyser was maintained throughout the duration of the study. The storage temperature of the samples were monitored and maintained throughout the study.

### Statistics

The median percentage difference of each parameter after each storage time at each temperature was calculated. This was compared to the maximum acceptable bias for that parameter as evidenced by biological variation studies^8,9^. A change or shift greater than the maximum acceptable bias was seen as a sign of unacceptable bias. The statistical significance of any such unacceptable bias was then tested by the Wilcoxon signed-rank test. The P-values of the Wilcoxon tests were adjusted for multiple comparisons by the Holm-Bonferroni method. All statistical tests were done using R statistical environment^10^.

For each parameter, the coefficient of variation calculated by the Root-Mean-Square Method at each temperature-time point of interest was also studied. The coefficients of variation (CV) at the different time-temperature points were compared against each other to see whether any marked increase occurred compared to the baseline at any time point. A rise in CV of 50% or more compared to the baseline CV constituted a marked shift which we deemed to be the appearance of instability in the automated reports.

## Results

The mean and standard deviations of the percent differences of each parameter after each time of storage at each temperature are given in Tables 1, 2 and 3.

**Table 1:**
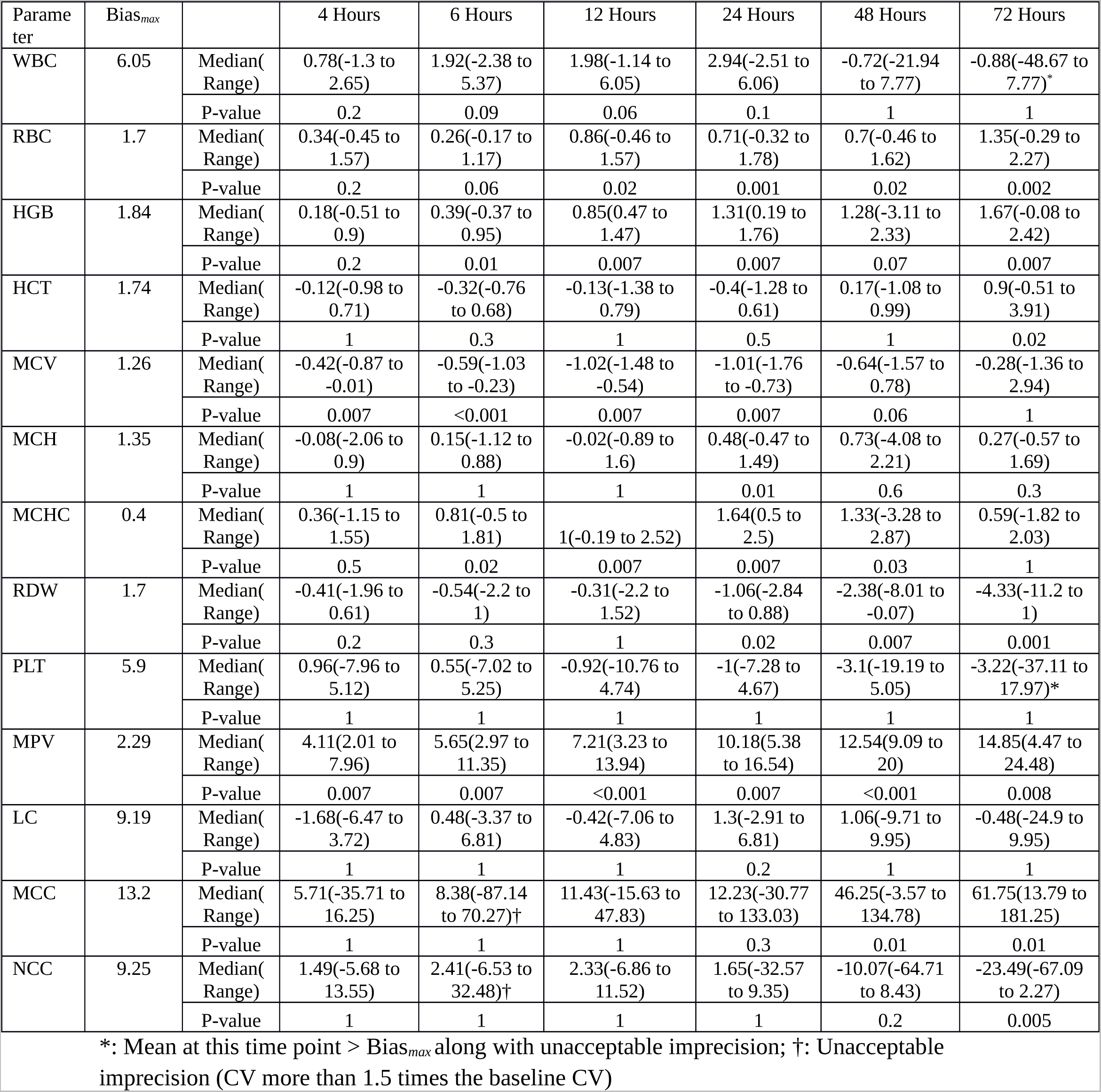
Showing the median, range of the Percent Difference of the studied parameters at different times of storage from the baseline, when stored at 4°C. The Holm-Bonferroni adjusted P-values of the Wilcoxon Signed Rank test are also given. (Bias*_max_*= Maximum acceptable Bias in %)

| Parameter | Bias<sub>max</sub> |  | 4 Hours | 6 Hours | 12 Hours | 24 Hours | 48 Hours | 72 Hours |
| --- | --- | --- | --- | --- | --- | --- | --- | --- |
| WBC | 6.05 | Median(Range) | 0.78(-1.3 to 2.65) | 1.92(-2.38 to 5.37) | 1.98(-1.14 to 6.05) | 2.94(-2.51 to 6.06) | -0.72(-21.94 to 7.77) | -0.88(-48.67 to 7.77)* |
|  |  | P-value | 0.2 | 0.09 | 0.06 | 0.1 | 1 | 1 |
| RBC | 1.7 | Median(Range) | 0.34(-0.45 to 1.57) | 0.26(-0.17 to 1.17) | 0.86(-0.46 to 1.57) | 0.71(-0.32 to 1.78) | 0.7(-0.46 to 1.62) | 1.35(-0.29 to 2.27) |
|  |  | P-value | 0.2 | 0.06 | 0.02 | 0.001 | 0.02 | 0.002 |
| HGB | 1.84 | Median(Range) | 0.18(-0.51 to 0.9) | 0.39(-0.37 to 0.95) | 0.85(0.47 to 1.47) | 1.31(0.19 to 1.76) | 1.28(-3.11 to 2.33) | 1.67(-0.08 to 2.42) |
|  |  | P-value | 0.2 | 0.01 | 0.007 | 0.007 | 0.07 | 0.007 |
| HCT | 1.74 | Median(Range) | -0.12(-0.98 to 0.71) | -0.32(-0.76 to 0.68) | -0.13(-1.38 to 0.79) | -0.4(-1.28 to 0.61) | 0.17(-1.08 to 0.99) | 0.9(-0.51 to 3.91) |
|  |  | P-value | 1 | 0.3 | 1 | 0.5 | 1 | 0.02 |
| MCV | 1.26 | Median(Range) | -0.42(-0.87 to -0.01) | -0.59(-1.03 to -0.23) | -1.02(-1.48 to -0.54) | -1.01(-1.76 to -0.73) | -0.64(-1.57 to 0.78) | -0.28(-1.36 to 2.94) |
|  |  | P-value | 0.007 | <0.001 | 0.007 | 0.007 | 0.06 | 1 |
| MCH | 1.35 | Median(Range) | -0.08(-2.06 to 0.9) | 0.15(-1.12 to 0.88) | -0.02(-0.89 to 1.6) | 0.48(-0.47 to 1.49) | 0.73(-4.08 to 2.21) | 0.27(-0.57 to 1.69) |
|  |  | P-value | 1 | 1 | 1 | 0.01 | 0.6 | 0.3 |
| MCHC | 0.4 | Median(Range) | 0.36(-1.15 to 1.55) | 0.81(-0.5 to 1.81) | 1(-0.19 to 2.52) | 1.64(0.5 to 2.5) | 1.33(-3.28 to 2.87) | 0.59(-1.82 to 2.03) |
|  |  | P-value | 0.5 | 0.02 | 0.007 | 0.007 | 0.03 | 1 |
| RDW | 1.7 | Median(Range) | -0.41(-1.96 to 0.61) | -0.54(-2.2 to 1) | -0.31(-2.2 to 1.52) | -1.06(-2.84 to 0.88) | -2.38(-8.01 to -0.07) | -4.33(-11.2 to 1) |
|  |  | P-value | 0.2 | 0.3 | 1 | 0.02 | 0.007 | 0.001 |
| PLT | 5.9 | Median(Range) | 0.96(-7.96 to 5.12) | 0.55(-7.02 to 5.25) | -0.92(-10.76 to 4.74) | -1(-7.28 to 4.67) | -3.1(-19.19 to 5.05) | -3.22(-37.11 to 17.97)* |
|  |  | P-value | 1 | 1 | 1 | 1 | 1 | 1 |
| MPV | 2.29 | Median(Range) | 4.11(2.01 to 7.96) | 5.65(2.97 to 11.35) | 7.21(3.23 to 13.94) | 10.18(5.38 to 16.54) | 12.54(9.09 to 20) | 14.85(4.47 to 24.48) |
|  |  | P-value | 0.007 | 0.007 | <0.001 | 0.007 | <0.001 | 0.008 |
| LC | 9.19 | Median(Range) | -1.68(-6.47 to 3.72) | 0.48(-3.37 to 6.81) | -0.42(-7.06 to 4.83) | 1.3(-2.91 to 6.81) | 1.06(-9.71 to 9.95) | -0.48(-24.9 to 9.95) |
|  |  | P-value | 1 | 1 | 1 | 0.2 | 1 | 1 |
| MCC | 13.2 | Median(Range) | 5.71(-35.71 to 16.25) | 8.38(-87.14 to 70.27)† | 11.43(-15.63 to 47.83) | 12.23(-30.77 to 133.03) | 46.25(-3.57 to 134.78) | 61.75(13.79 to 181.25) |
|  |  | P-value | 1 | 1 | 1 | 0.3 | 0.01 | 0.01 |
| NCC | 9.25 | Median(Range) | 1.49(-5.68 to 13.55) | 2.41(-6.53 to 32.48)† | 2.33(-6.86 to 11.52) | 1.65(-32.57 to 9.35) | -10.07(-64.71 to 8.43) | -23.49(-67.09 to 2.27) |
|  |  | P-value | 1 | 1 | 1 | 1 | 0.2 | 0.005 |

\*: Mean at this time point > Bias<sub>max</sub> along with unacceptable imprecision; †: Unacceptable imprecision (CV more than 1.5 times the baseline CV)

**Table 2:** Showing the median, range of the Percent Difference of the studied parameters at different times of storage from the baseline, when stored at 22°C. The Holm-Bonferroni adjusted P-values of the Wilcoxon Signed Rank test are also given. (Bias*_max_*= Maximum acceptable Bias in %)

| Parameter | Bias<sub>max</sub> |  | 4 Hours | 6 Hours | 12 Hours | 24 Hours | 48 Hours | 72 Hours |
| --- | --- | --- | --- | --- | --- | --- | --- | --- |
| WBC | 6.05 | Median(Range) | -0.08(-2.5 to 2.12) | 0.38(-1.33 to 5.71) | 0.44(-2.08 to 2.21) | 0.55(-3.8 to 4.35) | -2.34(-7.36 to 2.32) | -4.71(-10.93 to -0.16) |
|  |  | P-value | 1 | 1 | 1 | 1 | 0.1 | 0.006 |
| RBC | 1.7 | Median(Range) | 0.26(-1.55 to 1.44) | 0.18(-0.51 to 1.39) | 0.75(-1.08 to 2.09) | 0.85(-0.05 to 1.52) | 0.74(-0.91 to 1.32) | 0.48(-1.93 to 2.13) |
|  |  | P-value | 0.3 | 0.5 | 0.03 | 0.006 | 0.1 | 0.4 |
| HGB | 1.84 | Median(Range) | 0.15(-0.51 to 0.74) | 0.37(-0.07 to 1.16) | 0.8(0.31 to 1.64) | 1.08(0.19 to 1.75) | 1.32(0.32 to 2.63) | 1.5(-0.08 to 2.88) |
|  |  | P-value | 1 | 0.01 | 0.006 | 0.006 | 0.006 | 0.006 |
| HCT | 1.74 | Median(Range) | 0.62(-0.98 to 1.84) | 0.56(-0.84 to 1.69) | 1.29(-0.16 to 2.83) | 3.31(1.62 to 6.76) | 8.05(6.01 to 11.95) | 11.02(0.85 to 14.4) |
|  |  | P-value | 0.1 | 0.07 | 0.006 | 0.006 | 0.006 | 0.006 |
| MCV | 1.26 | Median(Range) | 0.24(-0.14 to 0.89) | 0.2(-0.38 to 1.21) | 0.78(-0.43 to 1.79) | 2.6(0.92 to 6.13) | 7.82(6.01 to 10.89) | 10.44(0.22 to 12.95) |
|  |  | P-value | 0.04 | 0.07 | 0.003 | <0.001 | <0.001 | <0.001 |
| MCH | 1.35 | Median(Range) | -0.15(-1.3 to 1.99) | 0.18(-1.19 to 1.04) | 0.15(-1.34 to 1.6) | 0.21(-1.19 to 1.24) | 0.75(-0.83 to 2.31) | 0.97(-0.65 to 3.14) |
|  |  | P-value | 1 | 1 | 1 | 0.3 | 0.3 | 0.01 |
| MCHC | 0.4 | Median(Range) | -0.64(-1.7 to 1.55) | -0.08(-2.04 to 0.96) | -0.64(-2.17 to 1.04) | -2.05(-5.61 to -0.46) | -6.46(-10.41 to -4.11) | -8.42(-11.39 to -0.93) |
|  |  | P-value | 1 | 1 | 0.3 | 0.006 | 0.006 | <0.001 |
| RDW | 1.7 | Median(Range) | 1.07(-0.31 to 2.92) | 1.41(-0.3 to 4.23) | 3.24(0 to 6.51) | 6.7(3.11 to 11.75) | 7.48(2.04 to 15.05) | 4.58(-0.9 to 19.36) |
|  |  | P-value | 0.007 | 0.007 | 0.007 | 0.006 | <0.001 | 0.01 |
| PLT | 5.9 | Median(Range) | 2.25(-4.12 to 5.03) | 1.61(-5.82 to 7.35) | 2.12(-3.35 to 6.4) | 2.4(-6.09 to 6.62) | 1.19(-9.78 to 8.93) | 1.23(-11.35 to 7.86) |
|  |  | P-value | 0.5 | 0.4 | 0.2 | 0.3 | 1 | 1 |
| MPV | 2.29 | Median(Range) | 7.62(2.66 to 10.42) | 7.8(1.7 to 10.9) | 7.39(2.11 to 12.5) | 8.23(1.51 to 13.65) | 11.85(6.82 to 19.43) | 12.63(7.27 to 22.2) |
|  |  | P-value | 0.006 | <0.001 | 0.007 | 0.006 | 0.006 | <0.001 |
| LC | 9.19 | Median(Range) | 0.73(-5.26 to 3.53) | 2.5(-3.37 to 9.22) | -0.97(-3.37 to 4.83) | 0.71(-6.47 to 4.83) | 3.54(-3.09 to 23.16) | 9.41(-4.2 to 34.95) |
|  |  | P-value | 1 | 0.5 | 1 | 1 | 0.1 | 0.001 |
| MCC | 13.2 | Median(Range) | 8.99(-6.25 to 79.55)† | 11.43(-0.83 to 43.48) | 22.65(8.14 to 56.52) | 5.9(-52.27 to 228.85) | -40.5(-61.36 to -19.64) | -63.13(-74.29 to 3.45) |
|  |  | P-value | 0.01 | 0.006 | 0.006 | 1 | NA | NA |
| NCC | 9.25 | Median(Range) | -2.13(-4.91 to 0.89) | -2.98(-9.8 to 0.73) | -3.15(-10.07 to -0.45) | -1.97(-31.92 to 14.99)† | 3.24(0.3 to 11.3) | -0.67(-8.61 to 10.57) |
|  |  | P-value | 0.07 | 0.01 | 0.006 | 1 | NA | NA |

†: Unacceptable imprecision (CV more than 1.5 times the baseline CV); NA: 10 or more invalid results

**Table 3:** Showing the median, range of the Percent Difference of the studied parameters at different times of storage from the baseline, when stored at 33°C. The Holm-Bonferroni adjusted P-values of the Wilcoxon Signed Rank test are also given. (Bias*_max_*= Maximum acceptable Bias in %)

| Parameter | Bias<sub>max</sub> |  | 4 Hours | 6 Hours | 12 Hours | 24 Hours | 48 Hours | 72 Hours |
| --- | --- | --- | --- | --- | --- | --- | --- | --- |
| WBC | 6.05 | Median(Range) | -0.06(-3.65 to 3.31) | -0.09(-3.75 to 2.62) | -1.11(-5 to 1.54) | -2.69(-8.75 to 0.71) | -5.56(-11.96 to -2.52) | -5.5(-15.94 to 1.79) |
|  |  | P-value | 1 | 1 | 0.02 | 0.004 | 0.004 | 0.001 |
| RBC | 1.7 | Median(Range) | 0.49(-0.16 to 2.04) | 0.54(-0.4 to 2.23) | 0.95(-0.4 to 2.12) | 0.9(-0.68 to 1.89) | 0.33(-1.87 to 1.39) | 0.34(-1.32 to 1.78) |
|  |  | P-value | 0.009 | 0.01 | 0.007 | 0.009 | 1 | 0.2 |
| HGB | 1.84 | Median(Range) | -0.04(-0.94 to 0.52) | 0.37(-0.37 to 1.28) | 0.63(0 to 1.23) | 0.94(0.44 to 2.3) | 1.57(0.38 to 3.01) | 2.11(-0.58 to 3.36) |
|  |  | P-value | 1 | 0.04 | 0.004 | 0.004 | 0.004 | 0.004 |
| HCT | 1.74 | Median(Range) | 1.72(1.06 to 3.89)* | 2.64(1.31 to 5) | 5.97(3.99 to 8.16) | 11.7(9.16 to 13.26) | 15.36(10.44 to 18.96) | 17.62(1.45 to 21.1) |
|  |  | P-value | <0.001 | 0.004 | <0.001 | <0.001 | <0.001 | <0.001 |
| MCV | 1.26 | Median(Range) | 1.3(0.75 to 2.22) | 2.17(1.51 to 2.82) | 5.28(3.35 to 6.81) | 10.6(8.35 to 12.41) | 15.11(11.37 to 17.81) | 16.66(1.02 to 20.46) |
|  |  | P-value | 0.004 | <0.001 | <0.001 | <0.001 | <0.001 | <0.001 |
| MCH | 1.35 | Median(Range) | -0.59(-2.4 to 0.68) | -0.28(-2.51 to 1.6) | -0.31(-1.93 to 0.78) | 0.22(-0.99 to 1.4) | 1.43(-0.41 to 2.79) | 2.05(-0.88 to 3.22) |
|  |  | P-value | 0.05 | 0.9 | 1 | 1 | 0.001 | 0.005 |
| MCHC | 0.4 | Median(Range) | -1.95(-4.64 to -0.66) | -2.21(-5.15 to -0.47) | -5.02(-7.47 to -3.03) | -9.39(-11.06 to -7.47) | -12.28(-13.9 to -7.89) | -12.53(-15.95 to -2.09) |
|  |  | P-value | 0.004 | <0.001 | 0.004 | 0.004 | 0.004 | <0.001 |
| RDW | 1.7 | Median(Range) | 2.53(0.83 to 3.81) | 3.62(0.86 to 5.95) | 6.66(3.93 to 11.32) | 9.4(3.9 to 15.62) | 8.05(1.59 to 19.36) | 11.39(2.05 to 28.67) |
|  |  | P-value | 0.004 | 0.004 | 0.004 | <0.001 | <0.001 | <0.001 |
| PLT | 5.9 | Median(Range) | -6.4(-28.72 to 0.39) | -7.23(-36.21 to 0.63) | -7.77(-39.89 to 0.32) | -6.24(-39.37 to 3.67) | -7.3(-35.82 to 1.58) | -4.73(-35.69 to 6.24) |
|  |  | P-value | <0.001 | 0.001 | <0.001 | 0.003 | 0.001 | 0.1 |
| MPV | 2.29 | Median(Range) | 6.33(2.52 to 15.77) | 4.51(0.9 to 13.02) | 4.45(-2.89 to 13.33) | 8.22(0.1 to 17.14) | 9.35(-0.08 to 17.59) | 6.35(-3.4 to 16.88) |
|  |  | P-value | 0.004 | <0.001 | 0.01 | 0.004 | <0.001 | 0.001 |
| LC | 9.19 | Median(Range) | 1.46(-3.09 to 7.77) | 2.32(-4.95 to 6.45) | 0.57(-4.85 to 12.87) | 1.47(-4.12 to 18.81) | 2.91(-5.71 to 22.51) | 8.18(-4.29 to 43.68) |
|  |  | P-value | 0.4 | 0.3 | 1 | 0.3 | 0.4 | 0.009 |
| MCC | 13.2 | Median(Range) | 13.04(1.65 to 32.86) | 15.48(-3.23 to 45.95) | 26.09(-2.33 to 63.04)† | -9.3(-86.36 to 233.87) | -62.02(-67.86 to 2.86) | 13.79(13.79 to 13.79) |
|  |  | P-value | 0.004 | 0.004 | 0.001 | 1 | NA | NA |
| NCC | 9.25 | Median(Range) | -3.82(-8.47 to 2.23) | -4.47(-10.78 to 0.89) | -9.09(-24.02 to 0.89)† | -5.42(-40.15 to 19.41) | -1.72(-9.24 to 5.14) | 1.14(1.14 to 1.14) |
|  |  | P-value | 0.009 | 0.005 | 0.005 | 0.3 | NA | NA |

\*: Mean percentage change at this time point is 2% which is greater than Bias<sub>max</sub> ; †: Unacceptable imprecision (CV more than 1.5 times the baseline CV); NA: 10 or more invalid results

RBC count was stable for seventy two hours irrespective of the temperature. HGB demonstrated an average shift of less than the maximum acceptable bias for at least forty eight hours irrespective of the storage temperature. WBC count and MCH showed acceptable bias for twenty four hours at 33°C and for at least forty eight hours at 22°C and 4°C. The percentage difference for PLT against the baseline showed unacceptable decrease of around 10% within four hours when stored at 33°C which did not further decrease appreciably. PLT did not exceed the maximum acceptable bias of 5.9% up to seventy two hours at 22°C and twenty four hours at 4°C.

MCV showed bias of greater than the maximum acceptable limit after twelve hours at 22°C and within four hours at 33°C, with the bias markedly increasing with time subsequently. The parameters related to Red Cell Volume, i.e. HCT, MCHC and RDW, show corresponding bias associated with storage time and temperature. Like MCV, HCT shows unacceptable bias after twelve hours when stored at 22°C and in less than four hours at 33°C. RDW shows unacceptable bias after twenty four hours when stored at 4°C, within six hours at 22°C and within four hours at 33°C. MCHC, being inversely related to MCV, showed a marked decrease with time at 22°C and above.

MPV showed a bias greater than the maximum acceptable bias at four hours at all the temperatures studied.

MCC showed unacceptable percentage difference from the baseline within six hours of storage at 22°C and within four hours of storage at 33°C. Also, MCC showed increased Coefficient of Variation at six hours when stored at 4°C, at four hours when stored at 22°C and beyond six hours when stored at 33°C. The coefficient of variation for NCC were increased beyond six hours under storage at 4°C and 33°C and beyond twelve hours under storage at 22°C. On storage for more than twenty four hours at 22°C and 33°C, ten or more readings did not give a valid value for MCC or NCC; these were taken as evidence for instability.

Other than the MCC and NCC, only in the case of WBC and PLT at seventy two hours when stored at 4°C, did the Coefficient of Variation show more than a 50% increase over the baseline value. The analytical precision was maintained for the other parameters at all the time points.

A post-hoc power analysis for the Wilcoxon signed rank test was done by the G*Power program^11^. To detect a bias equal to the maximum acceptable bias, and assuming the coefficient of variation of each parameter to be equal to that found in the present study, a sample size of twenty shows a power of greater than 90% for all parameters other than MCHC at a significance level of 0.00064(the Bonferroni corrected value) as well as a naïve significance level of 0.05. The power for detecting differences in MCHC is lower (approximately 76% at a naïve significance level of 0.05).

## Discussion

Analytical stability of haematological parameters after varying conditions of sample storage has been researched by many laboratory scientists^1–3,6,12–17^. However, we still have limited knowledge of analytical stability above 30°C. Only one study has reported on storage stability above 30°C, but only for storage up to twenty four hours. ^1^. A temperature of 33°C was close to the average high temperature during summer (April to July) at the place of analysis.

This study demonstrates that at 33°C, there may be unacceptable degradation of results for many of the parameters even in less than four hours. MCH, HGB, RBC counts, WBC counts and Lymphocyte Counts are the only parameters which are stable beyond six hours of storage at 33°C. The rapid appearance of bias at 33°C for most parameters underlines the need for proper storage and transport boxes in blood camps and extra-institutional sample collection centres. We also recommend that analysis be done as soon as possible. However, proper storage and transport boxes may not always be possible in a resource poor setting. Therefore, in case of receipt of samples exposed to temperatures of 33°C or more for more than four hours, we recommend that interpretation of the haemogram be done in a more cautious manner. In such conditions, the MCH may a better parameter for interpretation of anaemia in place of the MCV or MCHC. RDW and HCT also become unreliable markers in such conditions ^18^. For interpretation of the platelet count at 33°C, an initial dip of about 10% within the first few hours may be taken into account.

The three part differential count and MPV shows rapid instability under storage even under the more ideal conditions of storage at 4°C and 22°C. Rapid sequential increase in the MPV under storage is a known limitation of MPV estimation^19,20^, and may serve as a confounding factor in any prior or future studies involving these parameters. For the three part differential count, increase in imprecision is a significant problem that appears either before or near to the appearance of bias.

One limitation of the present study is that the same vial was repeatedly used for analysing the samples. This entailed bringing the samples stored under 4 °C and 33°C to room temperature at the time of analysis during the duration of the study. Such repeated “heat-cold” cycles may affect the results. The alternative lies in dividing the samples into different aliquots and only analysing one aliquot while keeping the other aliquots untouched. This would be more difficult since it requires a greater quantity of the blood sample with appropriate anti-coagulation. Also, aliquoting may itself be error prone due to variability arising from the lack of standardization of the aliquots for haematological testing. Moreover, the primary conclusion of the study (i.e. the appearance of unacceptable bias even before four hours for most of the parameters under storage at 33°C), remains unaffected by this limitation since the bias in these cases appeared before any “heat- cold “cycle. Similarly, the finding of acceptability of the bias for RBC, HGB, MCH and WBC for at least twenty four hours at 33°C should be robust since these parameters remain stable in spite of the potentially destabilizing influence of the “heat-cold” cycles.

Another potential limitation is that this study was done under controlled conditions with samples from a healthy population. The varying temperature in real life as well as confounding arising from improper draw, mixing, anticoagulation, or individual factors like presence of antibodies and activated complement may, conceivably, lead to different results. Studies done on patient samples as well as normal samples have shown few differences [15]; the differences are likely to be mild and should not affect the conclusions of the study.

The present study has to be interpreted keeping in mind that the analytical stability of a parameter depends on the analyser used^2^. The storage stability of parameters on other analysers using a different technique (e.g. Platelet Counts by the optical method) may vary.

This study may be extended by studying the stability of haematological parameters under even higher temperatures, as well as repeating the present study on 33°C on different analysers using different counting methodologies. Since most parameters were stable for less than four hours at 33°C, stability at 1 hour, 2 hour and 3 hours after collection should also be studied at 33°C and above. For generalizability, the present study should be replicated in different populations, and if necessary, with larger sample size.

In conclusion, the present study demonstrates acceptable bias of Haemoglobin, RBC counts, WBC counts, and MCH even at 33°C for at least twenty four hours after collection. It also demonstrates rapid appearance of bias (even before four hours), and therefore, unsuitability of reporting of most other parameters on sample storage at 33°C. The stability of most parameters (WBC, RBC, HGB, HCT, MCV, MCH, PLT and LC) for at least twenty four hours at 4 degrees centigrade, as well as the unacceptable bias of HCT and MCV after twelve hours at 22°C, is demonstrated.

## Acknowledgments

The Authors thank Transasia Biomedicals (Pvt) Ltd, India, for supplying the reagents for the tests free of cost.

## Notes

**Conflicts of Interest**: None declared

